# What explains the high island endemicity of Philippine *Rafflesia*? A species distribution modeling analysis of three threatened parasitic plant species and their hosts

**DOI:** 10.1101/2023.07.10.548437

**Authors:** Jasper J.A. Obico, R. Sedricke C. Lapuz, Julie F. Barcelona, Pieter B. Pelser

## Abstract

**Premise:** *Rafflesia* are rare holoparasitic plants. In the Philippines, all but one species are found only on single islands. This study aimed to better understand the factors contributing to this distribution pattern. Specifically, we sought to determine whether narrow environmental tolerances of host and/or parasite species might explain their island endemicity.

**Methods:** We used Maxent species distribution modeling to identify areas with suitable habitat for *R. lagascae*, *R. lobata*, and *R. speciosa* and their *Tetrastigma* host species. These analyses were carried out for current climate conditions as well as two future climate change scenarios.

**Key results:** Whereas species distribution models indicated suitable environmental conditions for the *Tetrastigma* host species in many parts of the Philippines, considerably fewer areas have suitable conditions for the three *Rafflesia* species. Some of these areas are found on islands from which they have not been reported. All three species will face significant threats as a result of climate change.

**Conclusions:** Our results suggest that limited inter-island dispersal abilities and/or specific environmental requirements are likely responsible for the current pattern of island endemicity of the three *Rafflesia* species, rather than the environmental requirements of their *Tetrastigma* host species.

---

Parasitic plants shape ecosystems. They often reduce the growth and reproduction of their host plants, and this can impact vegetation cycling and zonation, community biomass and structure, population dynamics, the diversity and behavior of herbivores, pollinators and seed dispersers, and even the abiotic features of their environment (Pennings and Callaway, 2002; Press and Phoenix, 2005; Watson et al., 2022). Parasitic plants are considered particularly vulnerable to the effects of habitat destruction and climate change due to their obligate dependence on their host plants (Mkala et al., 2022). Yet, little attention has thus far been given to examining how their distribution ranges and those of their hosts might change under different climate change scenarios (Zamora and Mellado, 2019; Cai et al., 2022; Mkala et al., 2022; Renjana et al., 2022). This might be a result of the negative stereotype associated with parasitism, especially for species that are, correctly or incorrectly, considered noxious weeds (Costea and Stefanović, 2009), but there are also other reasons. Accurately predicting distribution ranges is challenging for parasitic species for which host specificity information is sparse, because the distribution of parasites is limited by the geographic distribution and ecosystem requirements of their hosts (Costea and Stefanović, 2009; Cai et al., 2022). These same factors complicate effective conservation management of parasitic plants.

The Philippines is one of the centers of biodiversity of *Rafflesia* R.Br. (Brown, 1821; Rafflesiaceae), a Southeast Asian genus of obligate endo-holoparasitic plants growing in primary and secondary rain forests (Meijer, 1997; Nais, 2001; Barcelona et al., 2009). *Rafflesia* species have been referred to as the ‘giant pandas of the plant world’, given their rarity, bizarre endo-holoparasitic lifestyle, and large and foul-smelling flowers (Josephson, 2000) and their importance as icons of plant conservation. The Philippine archipelago is home to at least 14 of the 32 to 42 *Rafflesia* species currently recognized (Nickrent, 1997 onwards; Pelser et al. 2011 onwards; Galindon et al., 2016; Pelser et al., 2019; Siti-Munirah et al., 2021; Malabrigo et al., 2023). All are of conservation concern based on IUCN criteria (IUCN Standards and Petitions Committee 2019) or those used by the Philippine Department of Environment and Natural Resources (Pelser et al., 2011 onwards; Galindon et al., 2016; DENR, 2017; Malabrigo et al. 2023): four species are considered Critically Endangered, nine Endangered, and one Vulnerable. Thirteen of these are each only found on a single island. The only exception is *R. speciosa* Barcelona and Fernando (Barcelona and Fernando, 2002), which grows on Panay as well as on the nearby island of Negros (Pelser et al., 2019). Although *Rafflesia* are found in various parts of the Philippines, they are notably absent from many islands, including the sizable islands of Bohol, Cebu, Leyte, Mindoro, and Palawan (Fig. 1; Pelser et al., 2011 onwards). It is presently unclear why *Rafflesia* is not known from these islands, but perhaps they are present and remain to be discovered. Because they are endoparasites, *Rafflesia* plants can easily remain undetected when they are not in a reproductive state (e.g., Barcelona et al., 2007; Pelser et al., 2016; Renjana et al., 2022). However, it is also possible that their absence is a result of local extinction. This is plausible, because most Philippine *Rafflesia* species are only known from very few populations (Pelser et al., 2016, 2018) and even populations of species that are relatively common and widespread are often small and geographically and genetically isolated from each other (Pelser et al., 2017, 2018). Such populations may therefore be particularly vulnerable to habitat loss, degradation and fragmentation, which have impacted much of the natural forest ecosystems to which *Rafflesia* are confined (Mittermeier et al., 1998; Ong et al., 2002; Chokkalingam et al., 2006; Laurance, 2007; Barcelona et al., 2009).

**Figure 1.**
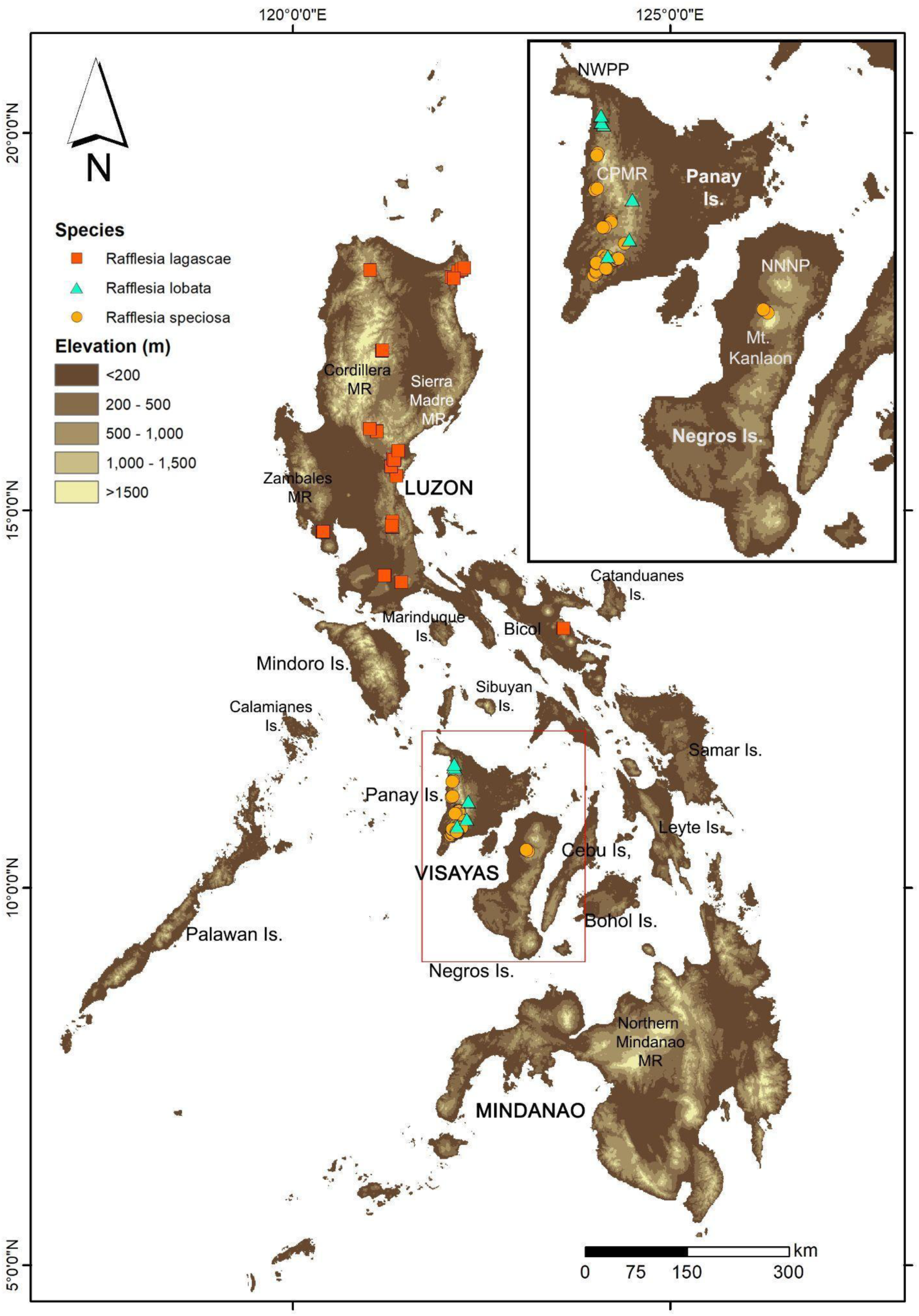
Map of the Philippines showing major island groups (in capital letters), islands (Is.), and mountain ranges (MR). Negros and Panay islands are highlighted within inset (boxed red in main map). Fill colors correspond to elevation in meters. *Rafflesia* occurrence points are overlaid to show native ranges of each species. CPMR = Central Panay Mountain Range; NNNP = Northern Negros Natural Park; NWPP = Northwest Panay Peninsula.

Pelser et al. (2019) suggested that if the high island endemism of *Rafflesia* is not a result of island-level extinction or a gap in our knowledge of their current distribution, it might be a result of poor inter-island dispersal ability, because the results of their molecular phylogenetic and biogeographic analyses indicate that dispersal of Philippine *Rafflesia* between islands has been relatively uncommon in their evolutionary history. Ants might be the primary seed dispersers of *Rafflesia* (Pelser et al., 2013, 2018). Because ants typically do not transport seeds further than 10–80 m (Gómez and Espadaler, 1998; Vittoz and Engler, 2008) and it can be assumed that large bodies of water are a significant barrier for myrmecochory, *Rafflesia* seeds might only be very rarely dispersed between islands.

It has been considered less likely that the absence of *Rafflesia* on some islands in the Philippines is due to the absence of suitable host species. Although *Rafflesia* species have only been observed to parasitize *Tetrastigma* (Miq.) Planch. (Planchon, 1887; Vitaceae) (Meijer, 1997; Nais, 2001; Pelser et al., 2016) and previous research suggests that Philippine *Rafflesia* might even display some level of host specificity or host preference at the species level, their reported hosts appear to be common and widespread in the Philippines (Pelser et al., 2016). Detailed information about the distribution and environmental requirements of these host species is however not currently available, because *Tetrastigma* are difficult to identify to species level due to their morphological plasticity and dioecious reproductive mode (Pelser et al., 2016; Obico et al., 2021a). Furthermore, specimens with reproductive parts (which can greatly assist species identification) can be difficult to detect in dense forest canopies and are therefore seldom collected (Obico et al., 2021a; Renjana et al., 2022).

*Rafflesia* species are not only strikingly absent from several large islands in the Philippines, but also from some areas within islands from which they have been reported (Pelser et al., 2018). These include areas that still have primary or relatively intact secondary forests and are within the elevational range throughout which *Rafflesia* have been found (i.e., c. 50–1380 m.a.s.l.; Barcelona et al., 2011; Pelser et al., 2018). For example, *Rafflesia* grow throughout the Central Panay Mountain Range (CPMR; Galang, 2009; Pelser et al., 2018), but they have never been reported from the forests of the nearby Northwest Panay Peninsula Natural Park (Fig. 1). Such observations and the disjunct distribution of populations of other Philippine *Rafflesia* species (e.g., Pelser et al., 2017) perhaps indicate that they have environmental requirements that are more specific than currently assumed (e.g., Pelser et al., 2018) or understood. Alternatively, or in addition to this, *Rafflesia* might not only disperse poorly between islands but also within islands, as could be inferred from studies that demonstrate strong genetic substructure within populations (Barkman et al., 2017) and poor genetic connectivity among populations that are more than c. 20 km away from each other (Pelser et al., 2018).

Here, we explore the hypothesis that Philippine *Rafflesia* species are range-restricted because of narrow environmental tolerances; either their innate tolerances or those of their *Tetrastigma* host species. We tested this hypothesis for three species of *Rafflesia* (*R. lagascae* Blanco (1845), *R. lobata* Galang and Madulid (2006), *R. speciosa*; Fig. 2) and their *Tetrastigma* hosts using a species distribution modeling approach. *Rafflesia speciosa* and *R. lobata* are currently classified as Endangered and *R. lagascae* is considered Vulnerable (Pelser et al., 2011 onwards; Malabrigo et al., 2023). A better understanding of differences between the observed and potential distribution patterns of *Rafflesia* species can inform their conservation management by identifying areas that have environmental conditions suitable for both parasite and host. These areas can then be prioritized when developing protected area networks (Ancheta et al., 2021; Renjana et al., 2022) or selected for establishing new *Rafflesia* populations when effective propagation and translocation techniques have been developed (Molina et al., 2017, 2023; Thorogood et al., 2022). Because of the potential negative impact of climate change on plant life (Kelly and Goulden, 2008), we further aim to contribute to the conservation of these three *Rafflesia* species by investigating projected future distribution patterns under two different climate change scenarios.

**Figure 2.**
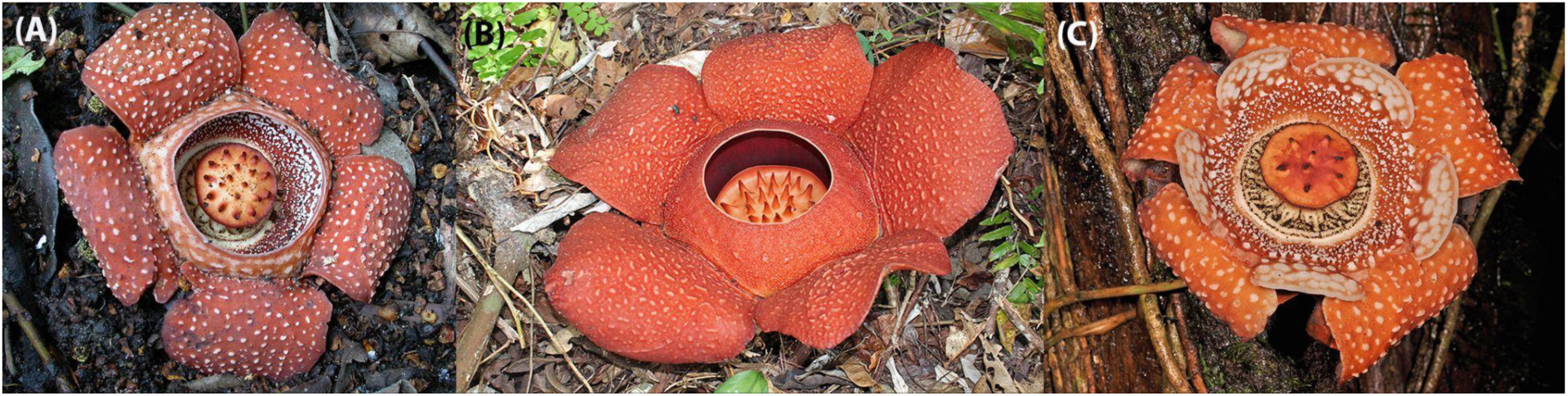
Flowers of (A) *Rafflesia lagascae*, (B) *R. speciosa*, and (C) *R. lobata*.

## MATERIALS AND METHODS

### Study species and distribution data

*Rafflesia lagascae* has relatively small flowers with a relatively wide diaphragm opening (Fig. 2). It is the most common *Rafflesia* species in the Philippines. It has the largest distribution area as well as the largest number of populations. *Rafflesia lagascae* is endemic to Luzon and grows from its northernmost provinces to the Bicol Peninsula in the south of the island (Fig. 1). It has a highly disjunct distribution pattern and most of the 11 populations that were included in a previous conservation genetic study showed poor genetic connectivity with other populations (Pelser et al., 2017). *Rafflesia lagascae* has been observed as a parasite of three *Tetrastigma* species: *T. ellipticum* Merr. (Merrill, 1916) s.l., *T. loheri* Gagnep. (Gagnepain, 1910), and *Tetrastigma* sp. A (sensu Pelser et al., 2016). All three are assumed to be common and have a wide distribution area in the Philippines, because they have been reported from all three major island groups of the Philippines: Luzon, Mindanao, and Visayas (Fig. 1). In addition to the *R. lagascae* populations referred to above, one other population (Mt. Labo in the Bicol Peninsula) has been identified as belonging to *R. lagascae*. Although the plants are morphologically similar to those of other populations and were resolved as closely related to them in a molecular phylogenetic study (Pelser et al., 2019), they are genetically distinct and may represent a cryptic species (Pelser et al., 2017). We therefore excluded locality records from the Mt. Labo population from the analyses included in the present study.

*Rafflesia speciosa* is a mid-sized *Rafflesia* with a relatively narrow diaphragm opening (Fig. 2). It has a much smaller distribution area than *R. lagascae*, but is locally common in the two areas from which it is known: the CPMR of the island of Panay and Mt. Kanlaon on Negros (Fig. 1). Pelser et al. (2018) observed little genetic structure among CPMR populations but substantial genetic differentiation between the CPMR and the single population known from Negros. *Tetrastigma harmandii* Planch. (Planchon 1887) and *T.* cf. *magnum* Merr. (Merrill, 1916; sensu Pelser et al., 2016) are the only known host species of *R. speciosa*. Like the aforementioned *Tetrastigma* hosts of *R. lagascae*, these two species have been found in Luzon, Mindanao and the Visayas.

*Rafflesia lobata* is morphologically similar to *R. lagascae* in producing small-sized flowers with a narrow diaphragm opening, but has a lobed diaphragm (Fig. 2) and is also phylogenetically distinct (Pelser et al., 2019). It is much rarer than *R. lagascae* and *R. speciosa*. *Rafflesia lobata* is restricted to the CPMR (Fig. 1) where it is only known from five populations, of which the southernmost three are possibly very small because we only observed one or two host plants for each. However, the two northernmost populations are comparatively large. Unpublished estimates of genetic connectivity among these populations using microsatellite data suggest limited gene flow between these two northern populations and the others (J. F. Barcelona and P. B. Pelser, unpublished data). *Rafflesia lobata* is sympatric with *R. speciosa* in a part of its known distribution area, but has a non-overlapping host range. However, its host species are the same as those of *R. lagascae*: *T. ellipticum* s.l., *T. loheri*, and *Tetrastigma* sp. A.

We recorded all occurrence points of these *Rafflesia* species and their *Tetrastigma* host species using Garmin GPS units during field work throughout the Philippines between 2007 and 2018 as part of our previous *Rafflesia* and *Tetrastigma* research projects (e.g., Barcelona, 2009; Pelser et al., 2013, 2016, 2017, 2018, 2019; Obico 2021a, b). Duplicates were removed and spatial thinning was implemented to retain only one occurrence point in every 30 arcsecond grid cell (i.e. the highest resolution at which bioclimatic data are available from CHELSA, see below; equivalent to ∼1 km^2^ at the equator) to avoid spatial sampling bias (i.e. unequal sampling effort resulting in a biased Maxent model; Kramer-Schadt et al., 2013). This resulted to 36 data points for *R. lagascae*, 6 for *R. lobata*, 40 for *R. speciosa*, 12 for *T. ellipticum* s.l., 24 for *T. harmandii*, 79 for *T. loheri*, 47 for *T.* cf. *magnum*, and 14 for *Tetrastigma* sp. A.

### Environmental variables for distribution modeling

Climate affects plant growth (Schimper, 1902), and the impacts of this become especially observable under the current accelerated climate change (Kelly and Goulden, 2008). However, edaphic factors also affect plant distribution (Witkamp, 1971) and should therefore also be included in species distribution models (SDMs) to assess whether potential changes in distribution are constrained by soil properties (Corlett and Tomlinson, 2020). To model the current distribution of *Rafflesia* and their hosts, both bioclimatic and soil variables were thus obtained from the CHELSA version 1.2 (Karger et al., 2017, 2018; https://chelsa-climate.org/bioclim/) and SoilGrids (Poggio et al., 2021; https://soilgrids.org/) databases, respectively. The bioclimatic data in CHELSA consist of a monthly temperature and precipitation climatology average for the years 1979–2013. Pearson’s correlation analyses were separately performed on the bioclimatic and soil variables to identify and exclude highly correlated (|r|> 0.7) variables (Pang et al., 2021), keeping variables that have been shown to be most important in explaining plant distribution patterns (Cai et al., 2003) or are predicted to undergo significant regional change in the future (Bindi et al., 2018; Philippine Atmospheric Geophysical and Astronomical Service Administration, 2018). The bioclimatic variables retained were mean annual air temperature (MAT; bio1), temperature seasonality (TS; bio4), annual precipitation (AP; bio12), precipitation seasonality (PS; bio15), and precipitation of the coldest quarter (PCQ; bio19), while the soil variables included bulk density, cation exchange capacity, soil pH, %clay, %silt, and %sand.

To constrain species modeling to the Philippines, bioclimatic and soil variables were cropped from the available global extent rasters using a shapefile of the Philippine archipelago. Soil variables were also resampled from their original 250 m resolution to 30 arcsecond using bilinear interpolation to make them compatible with the original resolution of the bioclimatic variables. All geoprocessing was implemented using the *raster* package (Hijmans, 2021) in R (R Core Team, 2021).

### Species distribution modeling

Species distribution models were generated for each individual *Rafflesia* and *Tetrastigma* species using Maxent (Phillips et al., 2006) implemented via the *dismo* package in R (Hijmans et al., 2020). For each species, six bootstrap replicates were run, and the jackknife setting was activated to determine individual variable contributions. This number is the lowest number of occurrences for any of the species included in our study (i.e., *R. lobata*). This is similar to the approach used by Banag et al. (2015). Earlier test runs which had more than six replicates did not improve model results significantly. The feature classes (FC) and regularization multipliers (RM) for each species were determined using the ENMevaluate() function of the *ENMeval* package in R (Kass et al., 2021). *ENMeval* examines a series of models across a range of settings for the FC and RM, and provides evaluation metrics to characterize model performance (Kass et al., 2021). In this study, the *ENMeval* analyses followed cross-validating model approaches by Pang et al. (2021): if species occurrence points were fewer or equal to 25, the ‘jackknife’ method was used; beyond 25, the ‘block’ method was applied. For each species, models with different combinations of FC (i.e., L, H, LQ, LQH, LQHP, and LQHPT; where L = linear, H = hinge, Q = quadratic, P = product, and T= threshold) and RMs (1 to 5, in increments of 0.5) were tested. A background of 10,000 points was used for all *Rafflesia* and *Tetrastigma* species. The model setting with the lowest *delta.AICc* value was ultimately selected as the most optimal.

Except for *T. harmandii* and *T. loheri*, the first set of SDMs generated contained some environmental variables with zero contribution. To further optimize the models, these variables were subsequently excluded, and Maxent analysis was repeated with only the remaining environmental variables and a different set of FC and RM. See Appendix S1 in Supplemental Data with this article for the Maxent settings used for each species. For the final SDMs, all five bioclimatic and soil variables were used, except for *R. lobata,* where only MAT and AP were retained.

To assess the reliability of the models, each SDM was evaluated using the Area Under the Curve (AUC) (Bradley, 1997) and the True Skill Statistic (TSS) (Allouche, 2006). Models were considered acceptable if they had both an AUC > 0.7 (Mandrekar, 2010) and a TSS value > 0.4 (Landis and Koch, 1977). Acceptable models were used to project the current and future species distributions onto space using the predict() function in *dismo*. The resulting continuous habitat suitability maps were then converted into binary maps using the minimum presence threshold (MPT) to distinguish areas with ‘suitable’ from those with ‘unsuitable’ habitat. The MPT, which was derived on the assumption that the least suitable habitat at which a species is known to occur is the species occurrence point with the lowest habitat suitability score (Pearson, 2007), ensures that all the *Rafflesia* and *Tetrastigma* species occurrence points are included in the projected suitable habitats for each species.

An unsuitability mask layer was created using a consensus land cover dataset generated by Tuanmu and Jetz (2014) to exclude areas without habitat that is assumed to be suitable for *Rafflesia*. The consensus land cover map that was used for this purpose is an integration of four land cover datasets (i.e., GlobCover, MODIS2005, GLC2000, DISCover) and was reported to have an improved ability to capture fine grain land cover data and accurately detect the presence of every class (Tuanmu and Jetz, 2014). Three land cover class layers that were considered unsuitable for *Rafflesia*, namely class 7 (cultivated and managed vegetation), class 9 (urban/built-up areas), and class 11 (barren land), were obtained from EarthEnv (https://www.earthenv.org/landcover; accessed on 14 June 2022) at 30 arcsecond resolution (Tuanmu and Jetz, 2014). To ensure occurrence points of *Rafflesia* and *Tetrastigma* species are not found within the boundaries of the unsuitable mask layer, the consensus prevalence, which is a measure of the overall agreement of the four integrated land cover datasets for a land cover class layer, was set to 100% for the three classes to completely exclude map cells classified under them. Furthermore, a raster file of the occurrence points of *Rafflesia* and *Tetrastigma* species was subtracted from the class 7 layer to exclude cells that were incorrectly classified as cultivated and managed vegetation based on our own observations at these field sites. The three land cover layers were then combined to generate the unsuitable mask layer, which was then applied onto the resulting SDMs. Instead of this post-hoc filtering approach, we could have directly included land cover data as one of the predictors in the SDMs, but that might have presented challenges for achieving model predictions with an appropriate level of accuracy, given that land cover is highly dynamic for Southeast Asia (Pang et al., 2021).

The percentages of suitable areas of each *Rafflesia* and *Tetrastigma* species in the Philippines were computed using the following formula: number of 30 arcsecond cells classified as ‘suitable’ divided by the total number of cells in the whole extent x 100. For the *Rafflesia* species, we also calculated the percentage of suitable area for each island (*R. lagascae, R. lobata*) or islands (*R. speciosa*) on which their presence has been confirmed in previous studies (hereafter referred to as ‘native island range’; Pelser et al., 2019).

To observe differences in environmental requirements among *Rafflesia* species and their respective *Tetrastigma* hosts, bioclimatic and soil values at predicted suitable areas were extracted per species using the extract() function of the *raster* package and displayed in box-and-whisker plots for native island ranges and the entire country.

### Future projections and change detection analyses

To ascertain the potential impact of climate change on species distributions, the future projections of the bioclimatic variables under two representative concentration pathways (RCPs), RCP 4.5 and RCP 8.5 for the years 2061–2080, were also obtained from CHELSA. The two RCPs represent the moderate and highest greenhouse gas emissions scenarios, respectively, according to the CMIP5 projections (van Vuuren et al., 2011). Three CMIP5 Earth System Models (ESMs) from CHELSA were used: CNRM-CM5, GFDL-CM3, and MPI-ESM-LR. These ESMs were reported to satisfactorily estimate Southeast Asian climate (McSweeney et al., 2015; Kamworapan and Surussavadee, 2019). The resulting SDMs from each ESM were averaged for each RCP scenario. Gain-loss statistics between current and future projections were then computed for each species using the Semi-automated Classification Plugin Tool in QGIS (Congedo, 2021). Grid cells that changed from unsuitable to suitable were counted as ‘gains’, while those that switched from suitable to unsuitable were recorded as ‘losses’. Cells that did not change were marked ‘stable’. Calculations were performed for both the native island ranges and the entire Philippines.

Suitable plant habitats have been found to shift to higher elevation as a result of global warming (e.g., Kelly and Goulden, 2008; Mancera and Lapuz, 2020). To detect such changes for our *Rafflesia* species under future climate scenarios, linear regression analyses were conducted between each species’ suitable habitat for each climate scenario and their respective elevations. We limited this analysis to the native island range since we are interested in potential changes to existing *Rafflesia* habitats. The projected suitable habitat rasters for each species were first converted from raster to occurrence point data using the as.data.frame() function in the *raster* package. These were then used to extract elevation data from the SRTM 90m Digital Elevation Data ver. 4.1 available from CGIAR-CSI (Jarvis et al., 2008; accessed on 5 Sep 2022) using the extract() function. The linear regression function lm() in the *stats* package in R was used separately for each *Rafflesia* species, with climate scenario and elevation (transformed to square root for normality) as the predictor and response variables, respectively. The p-values between groups were tested for significance using ANOVA type II, followed by Tukey’s HSD adjusted pairwise tests.

To observe differences in the edaphic characteristics between the suitable habitats under current and future climates, we also extracted edaphic data for the native island range of each species and tested differences in their values using the same method as outlined above.

### Protected area gap analysis

Lastly, suitable habitat area measurements were performed for each *Rafflesia* species to assess the protection that they receive under existing conservation regimes within their native island ranges. Protected Area boundary polygons for the Philippines were downloaded from the World Database on Protected Areas (UNEP-WCMC and IUCN, 2023). The polygons were clipped to restrict them to the native island range of each species, then overlaid on top of the suitable habitat rasters modeled under current conditions and future climate scenarios. The suitable habitat areas within and outside the polygons were measured for each climate scenario. The workflow utilized the *raster* and *tidyverse* (Wickham et al., 2019) packages in R (see Data Availability Statement).

## RESULTS

### Model accuracy

All species models for *Tetrastigma* and *Rafflesia* obtained acceptable average training AUC and TSS scores (Appendix S2). For *Tetrastigma* species, average training AUC scores ranged from 0.876–0.979, and average TSS scores ranged from 0.477–0.713. For *Rafflesia*, AUC values ranged from 0.970–0.997, and TSS scores ranged from 0.835–0.966. All species models were therefore considered for further analyses.

### Species distributions under current climate

#### Tetrastigma *hosts*

Because *Tetrastigma ellipticum* s.l.*, T. loheri*, and *Tetrastigma* sp. A are parasitized by *R. lobata* and *R. lagascae*, while *T. harmandii* and *T.* cf. *magnum* are parasitized by *R. speciosa*, the combined predicted suitable areas of each of these two host groups are presented separately. For both groups of *Tetrastigma* hosts, a large number of grid cells all over the Philippines were predicted to have suitable habitat (Fig. 3).

**Figure 3.**
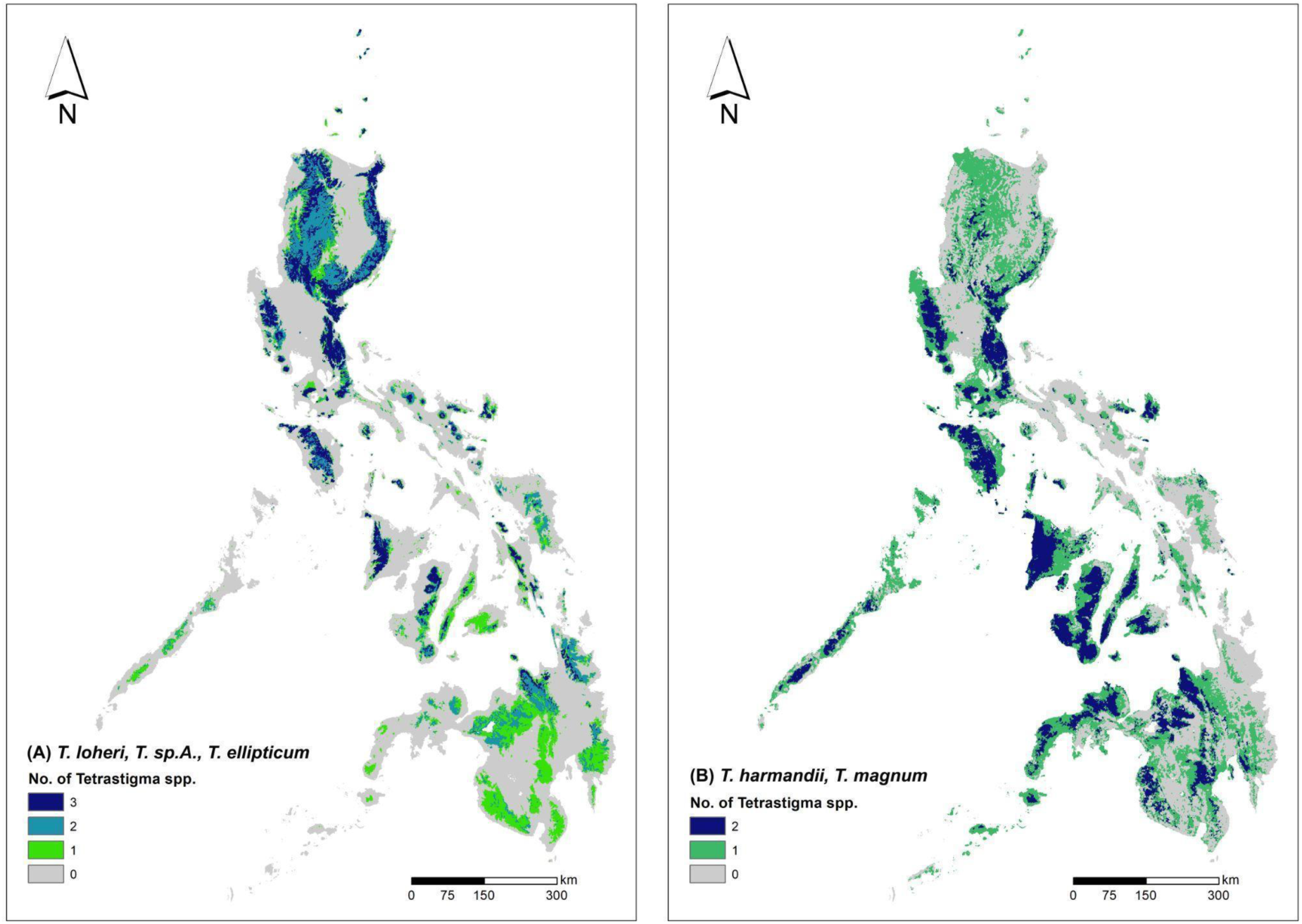
Distribution of suitable habitat of two *Tetrastigma* host groups: (A) the hosts of *R. lobata* and *R. lagascae* and (B) the hosts of *R. speciosa*. Colors show the number of *Tetrastigma* species that are predicted to occur in each grid cell.

#### Rafflesia *species*

Areas that are predicted to have suitable habitat for *R. lagascae* were found on several small and large islands in the Philippines, but mostly in the Luzon and central Visayas regions (Figs. 1 and 4). They cover 12.9% of the Philippine land area (Table 1). The largest portions of suitable habitat were found on mainland Luzon where its known populations are found. These represent 32.1% of the Luzon mainland area, and are concentrated in the island’s main mountain regions. Suitable habitat for *R. speciosa* was found in Negros and Panay, and in small areas in Calamianes and Mindoro (Figs. 1 and 5). The largest area with suitable habitat is on Panay Island. The suitable area covers 8.1% of Panay and Negros, which are the two islands from which *R. speciosa* has been reported. In total, only 0.7% of the Philippines provides suitable habitat for this species (Table 1). Areas with suitable habitat for *R. lobata* can be found on large and small islands across the Luzon, Visayas, and Mindanao regions (Figs. 1 and 6), and these cover 5.9% of the Philippines. On Panay Island where *R. lobata* is an endemic species, the suitable habitat covers 7.7% of the island and is mostly located in the CPMR.

**Figure 4.**
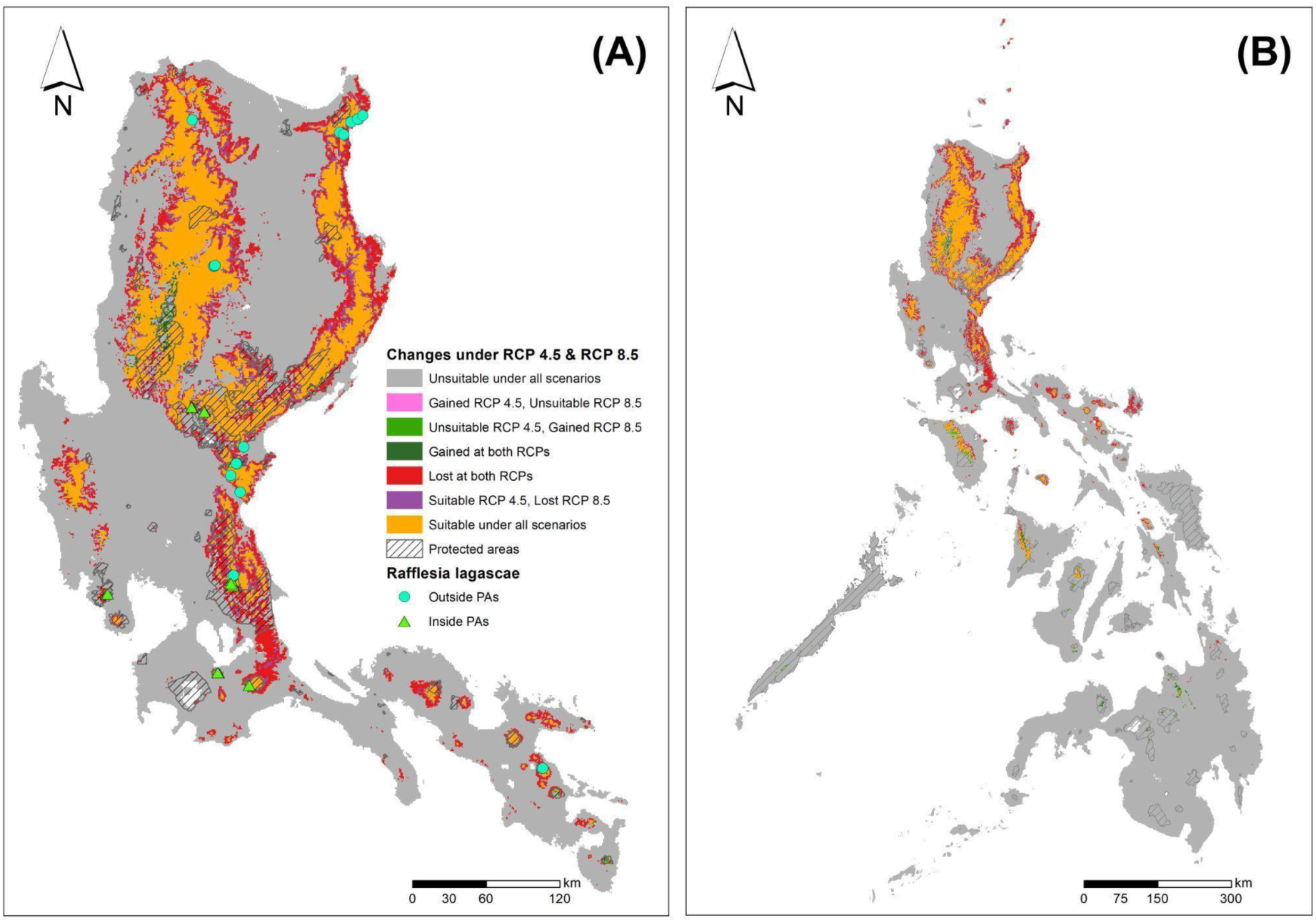
Distribution of suitable habitat of *Rafflesia lagascae* for (A) Luzon, the only island from which it is known and (B) the entire Philippines and combined changes to this under RCP 4.5 and RCP 8.5 from current climate. ‘Suitable under all scenarios’ refers to consistently suitable habitats from current to future climate scenarios. ‘Unsuitable under all scenarios’ refers to consistently unsuitable habitats from current to future climate scenarios. ‘Gained’ refers to habitats that became suitable under future climate, while ‘Lost’ refers to habitats that became unsuitable under future climate. Hatched areas are protected areas. Species points are labeled based on their inclusion within protected area boundaries. Calculated with *Rafflesia* SDMs.

**Figure 5.**
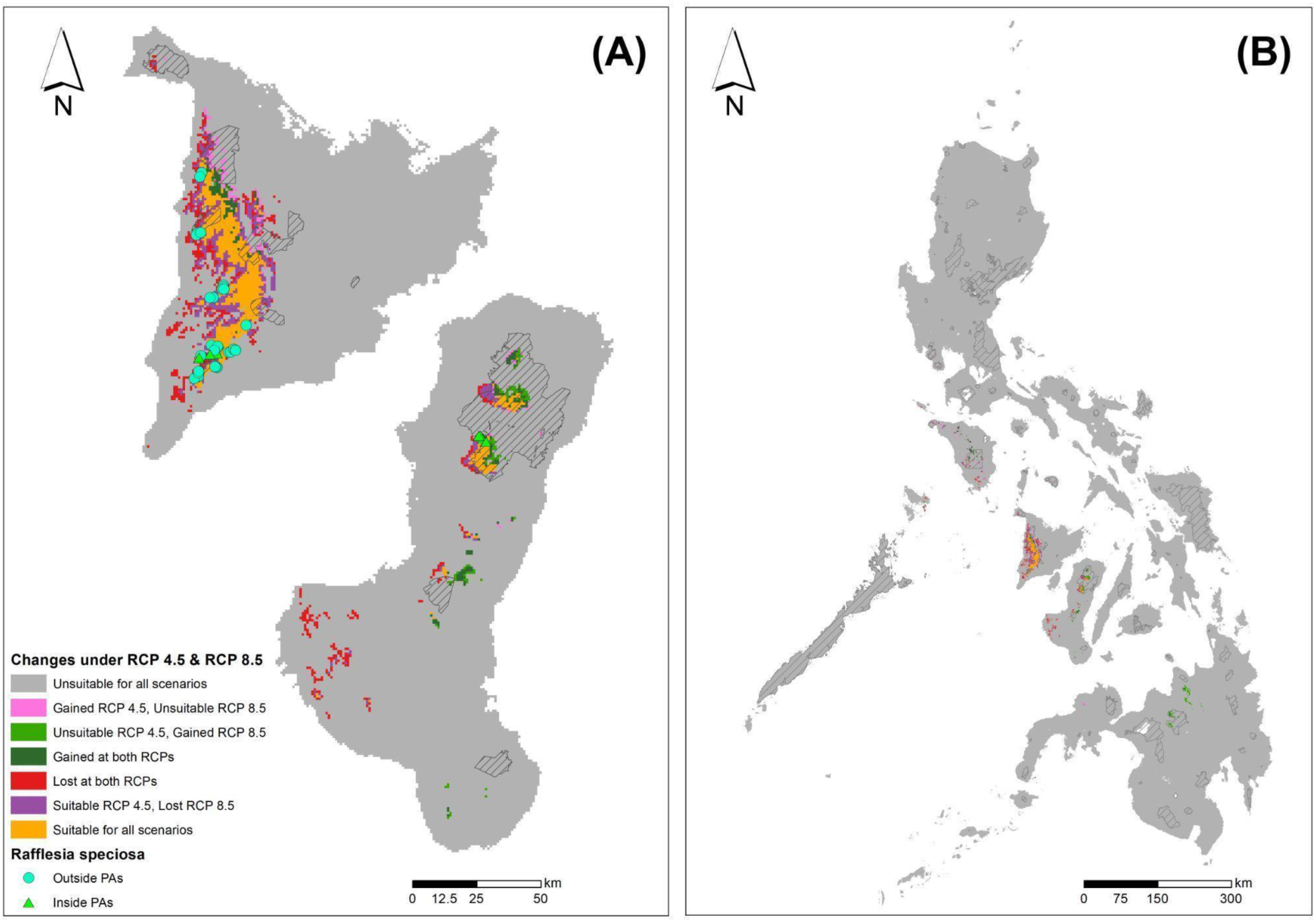
Distribution of suitable habitat of *Rafflesia speciosa* on (A) Panay and Negros, the only islands from which it is known and (B) the entire Philippines and combined changes to this under RCP 4.5 and RCP 8.5 from current climate. ‘Suitable under all scenarios’ refers to consistently suitable habitats from current to future climate scenarios. ‘Unsuitable under all scenarios’ refers to consistently unsuitable habitats from current to future climate scenarios ‘Gained’ refers to habitats that became suitable under future climate, while ‘Lost’ refers to habitats that became unsuitable under future climate. Hatched areas are protected areas. Species points are labeled based on their inclusion within protected area boundaries. Calculated with *Rafflesia* SDMs.

**Figure 6.**
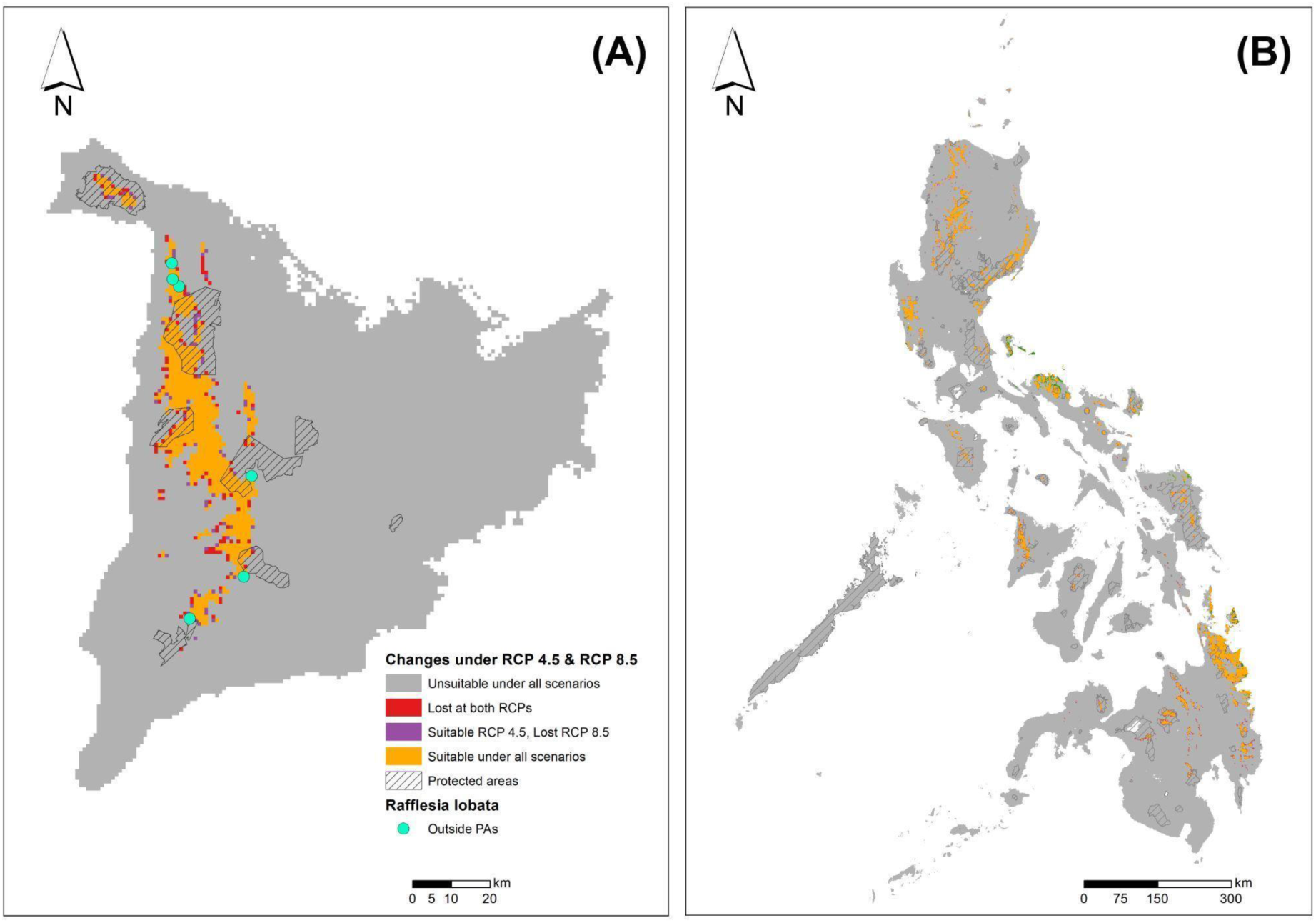
Distribution of suitable habitat of *Rafflesia lobata* on (A) Panay, the only island from which it is known and (B) the entire Philippines and combined changes to this under RCP 4.5 and RCP 8.5 from current climate. ‘Suitable under all scenarios’ refers to consistently suitable habitats from current to future climate scenarios. ‘Unsuitable under all scenarios’ refers to consistently unsuitable habitats from current to future climate scenarios. ‘Gained’ refers to habitats that became suitable under future climate, while ‘Lost’ refers to habitats that became unsuitable under future climate. Hatched areas are protected areas. Species points are labeled based on their inclusion within protected area boundaries. Calculated with *Rafflesia* SDMs.

**Table 1.**
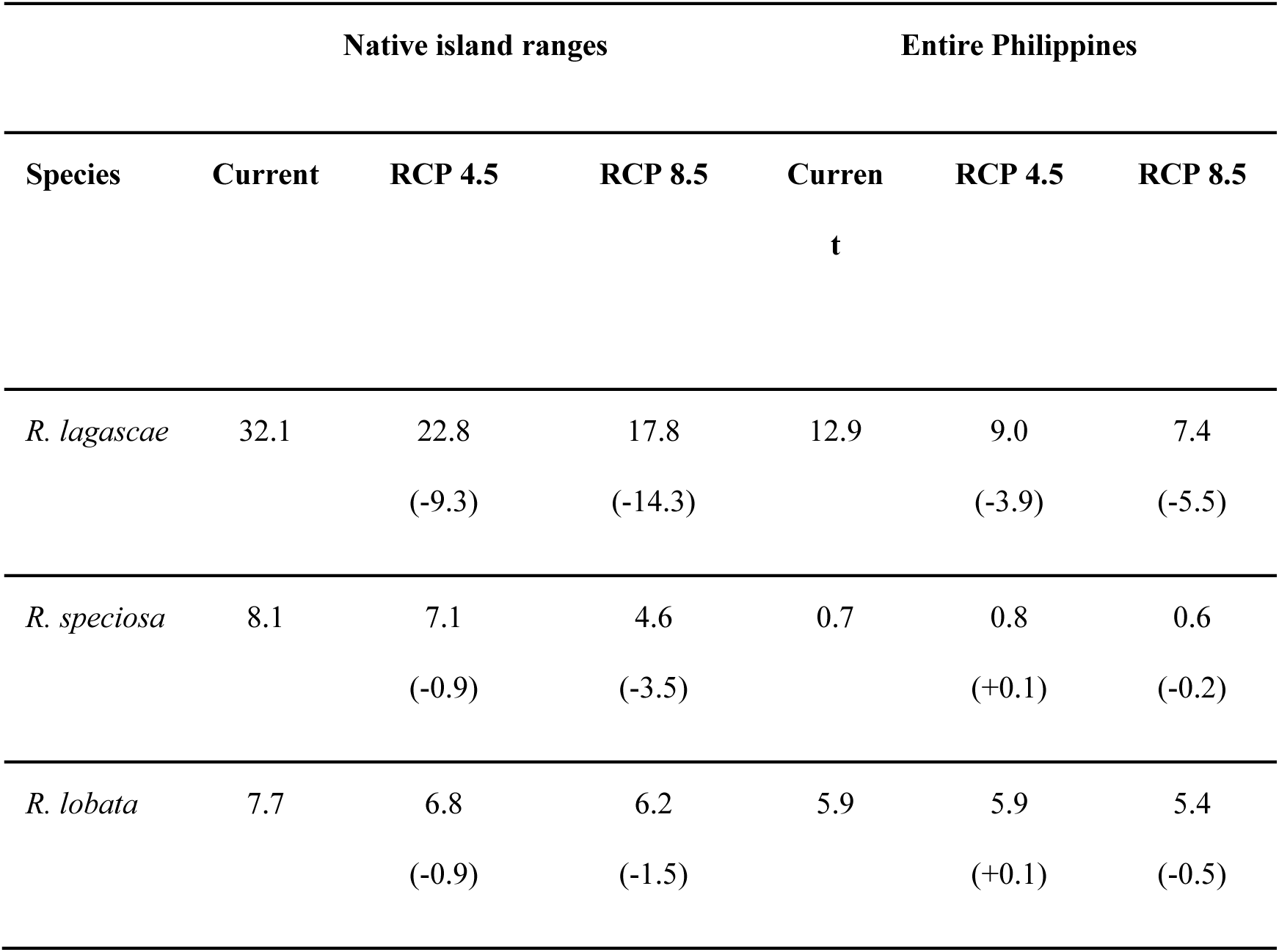
Percentages of areas with suitable *Rafflesia* habitat for the native island ranges and the entire Philippines under current climate conditions and two future climate change scenarios (i.e., RCP 4.5, RCP 8.5) resulting from *Rafflesia* SDMs. Values in brackets indicate the amount of change for that climate scenario when compared against predictions under the current climate.

### Environmental requirements of Rafflesia species

The most important variables contributing to the SDM of *R. lagascae* were climate variables, of which mean annual temperature contributed most to the model (45.5%) (Appendix S3). Soil pH had a contribution of 11%. The climate variables were also the top contributors to the model for *R. speciosa* and had a total contribution of 87.3% with precipitation seasonality as the highest contributor (23.2%). The soil variables had a relatively low contribution of only 12.9% (Appendix S3). The prediction of the suitable habitat for *R. lobata* was only based on the contribution of two climate variables: annual precipitation (70.1%) and mean annual temperature (29.9%), and no soil variables were included (Appendix S3).

Figure 7 summarizes the relative differences in environmental requirements among the three *Rafflesia* species. These results indicate that *R. speciosa* occupies the warmest niches and *R. lobata* the coolest. *Rafflesia lobata* grows in the wettest environments and *R. lagascae* prefers the least precipitation. *Rafflesia speciosa* habitats have the narrowest range of temperature seasonality but the highest precipitation seasonality. The three species are also different in their soil requirements. *Rafflesia speciosa* grows in hosts that prefer more basic soils. Of the three species, its host plants grow on the sandiest soils that have the least clay. On the other hand, *R. lagascae* host plants prefer soils with higher silt and clay content. The largest differences between hosts and their parasites in their suitable environmental ranges were observed for *R. speciosa* (Appendix S4). This species showed considerably narrower ranges than its hosts in all environmental variables included in our analyses.

**Figure 7.**
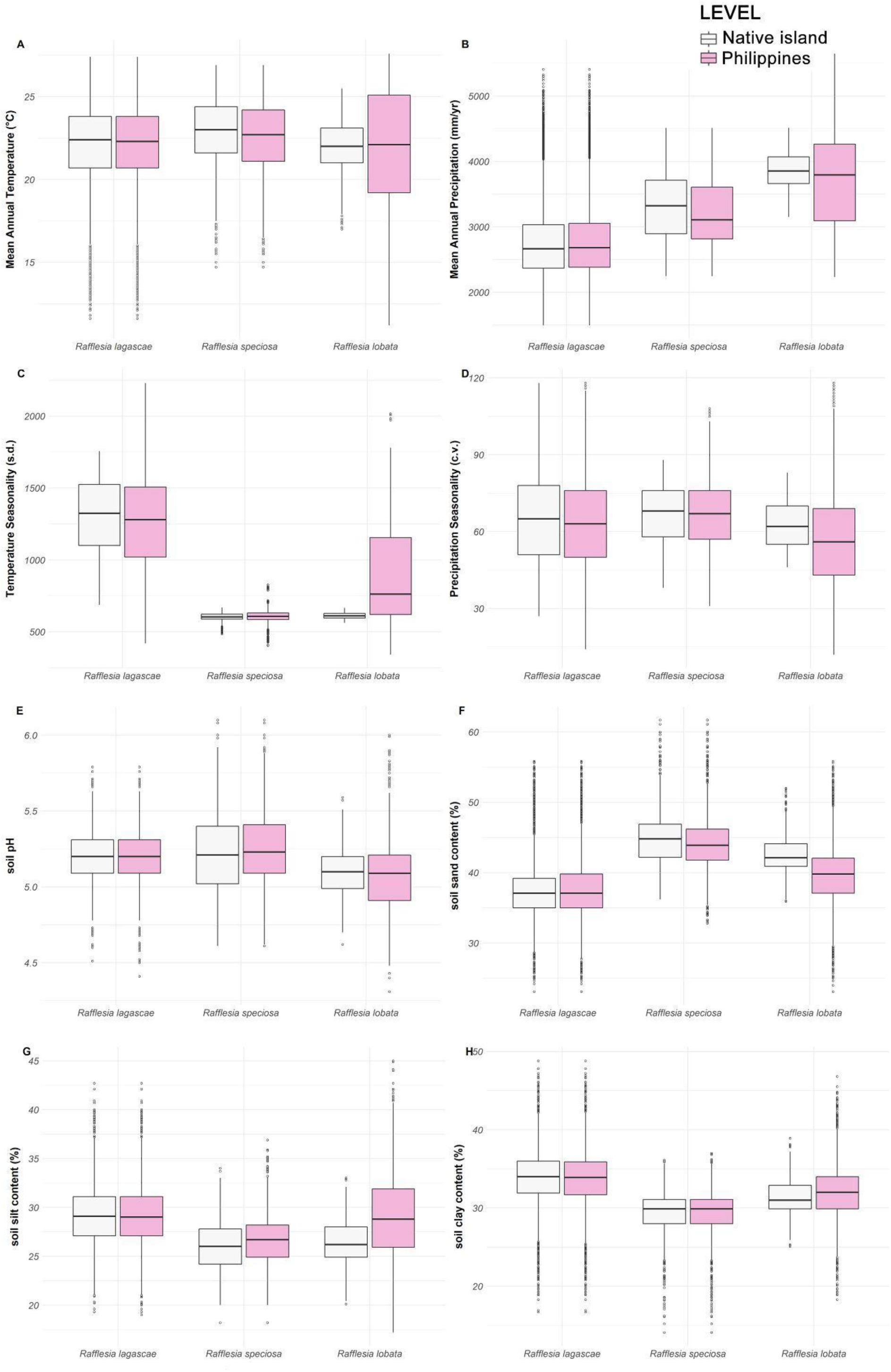
Boxplots depicting the differences in environmental requirements among *Rafflesia* species (s.d.: standard deviation; c.v.: coefficient of variation) estimated for the islands on which each species has been found (native island, white boxes) and the entire Philippines (pink boxes). Calculated with *Rafflesia* SDMs.

### Future projected distribution changes in *Rafflesia*

All modeled *Rafflesia* species are generally predicted to lose suitable habitat under both future climate scenarios, with higher losses observed under RCP 8.5 (Table 1 and Appendix S5; Figs. 4–6). *Rafflesia lagascae* will suffer the greatest losses within the currently known distribution areas among the three: its total percentage of area in Luzon with suitable habitat might be reduced by 9.3% (RCP 4.5) or 14.3% (RCP 8.5) (Table 1). Most of the losses are observed along the fringes of the current area of suitable habitat, and are more pronounced along the southern portions of the Sierra Madre Mountain Range and Bicol peninsula (Figs. 1 and 4). When areas with suitable environmental conditions are considered that are also elsewhere in the Philippines, the total area declines by 3.9% under RCP 4.5 and 5.5% under RCP 8.5 (Table 1), which signifies predicted losses of suitable habitat outside the present range of *R. lagascae*. Minor gains (<1% of suitable habitat) are observed in the central portions of the Cordillera mountain range in Luzon, and also in central portions of the mountain ranges of Mindoro, Panay, and Northern Mindanao (Figs. 1 and 4).

The total area with suitable habitat for *R. speciosa* in Negros and Panay is projected to decline by 0.9% under RCP 4.5 and 3.5% under RCP 8.5 (Table 1). Most of these losses are predicted for Panay Island (Fig. 5). When considering the whole of the Philippines, there are negligible gains predicted under RCP 4.5. Losses are observed under RCP 8.5 (0.2%; Table 1), despite a potential gain of new suitable areas in northern Mindanao (Fig. 5).

Among the three species, *R. lobata* might be the least affected by climate change. The area with suitable habitat in Panay is only predicted to decline by 0.9% (RCP 4.5) and 1.5% (RCP 8.5; Table 1). The losses are all predicted to occur along the margins of its suitable areas in Panay (Fig. 6). Not only is the amount of suitable habitat predicted to decrease for all three species, areas with such habitat are also predicted to shift to higher elevations on the islands where the three *Rafflesia* species are found (Fig. 8). For each species, a significant (*R. lagascae*: F(2.92) = 5154.76, p<0.001; *R. speciosa*: F(2.56) = 641.85, p<0.001; *R. lobata*: F(2.28) =12.08, p<0.001) increase in elevation is expected in under both climate change scenarios (RCP 4.5: 180 m, RCP 8.5: 291 m for *R. lagascae*; RCP 4.5: 35 m, RCP 8.5: 57 m for *R. lobata*; RCP 4.5: 153 m, RCP 8.5: 349 m for *R. speciosa*; Appendix S6). Climate change is also expected to result in changes in edaphic variables within the suitable habitats of the *Rafflesia* species under both climate change scenarios (Appendix S7). Although these changes were small, they were statistically significant (p<0.001) for most variables.

**Figure 8.**
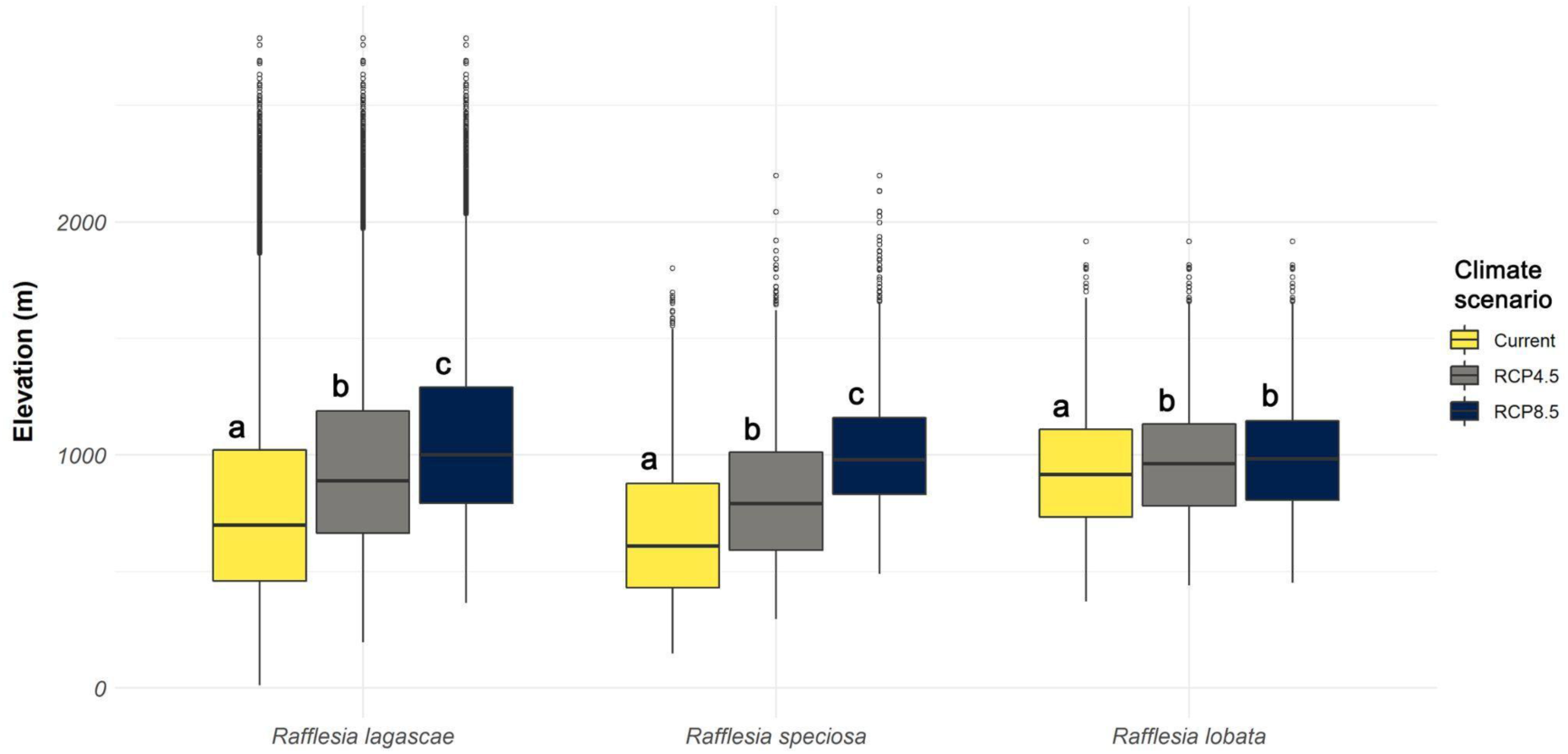
Boxplot showing elevational changes in suitable habitats of *Rafflesia* species under current and future climate scenarios (RCP 4.5 and RCP 8.5) on islands on which they are currently present. Letters indicate differences among climate scenarios that are statistically significant at p <0.05 as tested using ANOVA type II, followed by Tukey’s HSD adjusted pairwise tests. Calculated with *Rafflesia* SDMs.

### Protected area gaps

The suitable habitat within protected areas under current climate for *Rafflesia lagascae,* which has the largest percentage at 68.0% (8369 km²), is projected to decrease to 50.8% under RCP 4.5 and 40.6% under RCP 8.5 (Table 2). *Rafflesia lobata* is likewise projected to lose suitable habitat within its native island range: from 31.4% under current climate to 27.4% (RCP 4.5) and 24.5% (RCP8.5). For *R. speciosa* which has the smallest protected suitable area (440 km²), a slight increase is projected under RCP 4.5 (from 17.0% to 20.4%), but a decrease under RCP 8.5 (to 15.6%).

**Table 2.** Areas (km^2^) and percentages of suitable habitats within protected area boundaries for *Rafflesia* species in their native islands. Calculated with *Rafflesia* SDMs.

| Species | Current | RCP 4.5 | RCP 8.5 |
| --- | --- | --- | --- |
| *R. lagascae* | 8369 / 68.0% | 6253 / 50.8% | 5001 / 40.6% |
| *R. speciosa* | 440 / 17.0% | 530 / 20.4% | 406 / 15.6% |
| *R. lobata* | 273 / 31.4% | 238 / 27.4% | 213 / 24.5% |

## DISCUSSION

### What underlies the restricted distribution of Philippine Rafflesia?

This study sought to better understand the current distribution patterns of three *Rafflesia* species endemic to the Philippines using species distribution models for these parasitic plants and their *Tetrastigma* host species. Its main aim was to determine if narrow environmental tolerances of *Tetrastigma* and/or *Rafflesia* species might be contributing factors to the limited distribution of the latter, with *R. lagascae* and *R. lobata* confined to one island and *R. speciosa* to two. Addressing this question will improve our understanding of the impact of climate change on parasitic plants. This is important because this is presently very poorly known, despite the importance of parasitic plants to community-level ecological interactions (Press & Phoenix, 2005; Zamora and Mellado, 2019; Mkala et al., 2022; Watson et al., 2022).

The SDM results indicate that suitable environmental conditions for all five *Tetrastigma* species included in this study are present on all or most large islands of the Philippines (Fig. 3). SDM maps produced for both *Rafflesia* host groups (i.e., *T. ellipticum* s.l., *T. loheri*, and *T.* sp. A. for *R. lagascae* and *R. lobata*; *T. harmandii* and *T.* cf. *magnum* for *R. speciosa*) further suggest that five large islands from which *Rafflesia* species have never been recorded (i.e., Bohol, Cebu, Leyte, Mindoro, Palawan) provide considerable areas of suitable habitat for at least one of the observed host species for each of the three *Rafflesia* species (Fig. 3). This suggests that it is unlikely that *R. lagascae, R. lobata* and *R. speciosa* are absent from these islands because they do not provide a suitable environment for the hosts of these *Rafflesia* species. Although it is technically possible that *Tetrastigma* hosts might be absent from an island for other reasons than the absence of suitable habitat, which then could explain the absence of *Rafflesia*, we consider this an unlikely general explanation for most, if not all, five islands. Empirical data show that all five species are widespread in the Philippines (Pelser et al., 2011 onwards, 2016; Obico et al., 2021a, b). For example, *T. loheri* has been reported from Cebu, Leyte, Mindoro, and Palawan (Pelser et al., 2011 onwards).

These results provide support for the hypothesis that the environmental requirements of *Tetrastigma* species are not a limiting factor to the current distribution of the *Rafflesia* species included in our study (Pelser et al., 2018). This pattern supports the ‘parasite niche hypothesis’ (Lira-Noriega and Peterson, 2014): the distribution of parasites is mediated by their autecology instead of the distribution of their hosts. Evidence in support of this hypothesis has been found for mistletoe species in families Loranthaceae and Viscaceae (Lira-Noriega and Peterson, 2014; Ramírez-Barahona et al., 2017; Ornelas et al., 2018).

If host distribution is not a limiting factor to the distribution of *R. lagascae*, *R. lobata* and *R. speciosa* under current climate conditions, it is possible that these species have relatively few populations and small observed distribution areas because they have quite specific environmental requirements. If this is the primary reason for this distribution pattern, then we expect that their recorded range would closely correspond to the areas with suitable habitat as identified in the SDM analyses. However, this is not the case for all three *Rafflesia* species.

*Rafflesia speciosa* seems to have the most restrictive habitat requirements of the three species. The SDM results indicate that only 0.7% of the terrestrial area of the Philippines provides suitable habitat for this species (Table 1). A total of 91.2% of this area is found within the two islands that compose its known distribution area (i.e., Panay and Negros; Fig. 5), where 8.1% of their land mass was identified as having suitable conditions for *R. speciosa* (Table 1). Suitable habitat was also predicted for other islands (i.e., Calamianes and Mindoro), but only in relatively small and disjunct areas (Fig. 5). Even within Panay and Negros, the most contiguous areas with suitable habitat are those from which this species is currently known (Fig. 5). Although an area in Northern Negros Natural Park north of the Mt. Kanlaon population of *R. speciosa* seems to be the exception to this pattern, anecdotal reports suggests that the area might in fact be home to a second Negros population of this species (S. Olimpos, personal communication). Therefore, *R. speciosa* might primarily be confined to its currently known distribution area because of narrow environmental tolerances The results of our SDM analyses suggest that the relatively. small area in the Philippines that has suitable habitat for *R. speciosa* compared to *R. lagascae* and *R. lobata* could largely be a consequence of the combination of its much narrower tolerance to changes in temperature seasonality (Fig. 7c) and its preference for higher precipitation seasonality (Fig. 7d).

*Rafflesia speciosa* shares Panay with *R. lobata*, which is endemic to the CPMR. In contrast to *R. speciosa*, sizable areas with suitable habitat for *R. lobata* might be present on several islands from which this species has never been reported, including the mountain ranges of Luzon, northern and eastern Mindanao, Mindoro, and Samar (Fig. 6), despite presence of their *Tetrastigma* hosts (Pelser et al., 2011 onwards, 2016; Obico 2021a). In addition, whereas suitable habitat for *R. speciosa* does not appear to be present in the Northwest Panay Peninsula, it may have suitable habitat for *R. lobata*, although this species has not been reported from this area notwithstanding its proximity to the CPMR and presence of *T. loheri* (J. F. B. & P. B. P., pers. obs.) on the peninsula. *Rafflesia lobata* is the only one of the three *Rafflesia* species for which the largest proportion of its suitable habitat is found outside of the islands to which it is restricted (94.9%, vs. 8.9% for *R. speciosa* and 9.5% for *R. lagascae*). These SDM patterns indicate that narrow environmental tolerances do not adequately explain the island endemicity of *R. lobata* and that this may instead be a result of poor inter-island dispersal abilities, as has been previously suggested (Pelser et al., 2019).

Suitable habitat for *R. lagascae* seems to be present throughout the Cordillera, Sierra Madre and Zambales mountain ranges of Luzon, as well as in the mountains of the Bicol peninsula (Figs. 1 and 4). These are all areas from which this species has previously been reported. However, although these areas comprise most of the areas with suitable habitat for *R. lagascae* in the Philippines (90.5%), the results of the SDM analyses indicate that sizable areas that provide appropriate environmental conditions for this species and that have forests from which *Tetrastigma* host species have been recorded (Pelser et al., 2011 onwards) can also be found on islands in the central part of the Philippines, most notably Catanduanes, Marinduque, Mindoro, Negros, Panay and Sibuyan (Figs. 1 and 4). Thus, *R. lagascae* displays SDM patterns that are somewhat intermediate between those of *R. lobata* and *R. speciosa* and its endemicity to Luzon might be explained by a combination of limited inter-island dispersal ability and environmental conditions that are not present on some major islands in the Philippines.

Climate, soil, and elevation data in the observed *Rafflesia* populations and modeled with Maxent provided details about the different habitat requirements of the three species. These show differences in temperature, precipitation, temperature and precipitation seasonality, and soil composition requirements among the species (Fig. 7). *Rafflesia speciosa* and *R. lobata* both grow in the CPMR, but are only found in sympatry at two locations. This might be explained by their quite different environmental needs. *Rafflesia speciosa* grows in the parts of the mountain range that are warmer and drier and that experience more precipitation seasonality than the areas where *R. lobata* can be found (Fig. 7). Its hosts prefer soils that are more basic and have a lower clay content. These sites are commonly found at a lower elevation than those of *R. lobata* (Fig. 8). These different environmental requirements and their different flower sizes (11–21 cm diameter for *R. lobata* vs. 45–56 cm diameter for *R. speciosa*; Barcelona et al., 2011) might indicate the importance of divergent selection in the evolutionary history of *Rafflesia* (Pelser et al., 2019). This might also be suggested by their non-overlapping host preferences (Pelser et al. 2016), although the latter could also be a result of competitive exclusion. SDM and other studies that provide information about the environmental needs of other *Rafflesia* species (e.g., Manuel and Hermocilla, 2021) that live in the same areas but that are not or only partially sympatric could provide additional insights into the role of divergent selection through habitat specialization. *Rafflesia lagascae* could be a useful focal species for such studies in the Philippines because it is partially sympatric with *R. consueloae* Galindon, Ong and Fernando, *R. leonardi* Barcelona and Pelser, and *R. philippensis* Blanco (Blanco, 1845; Barcelona et al., 2008; Pelser et al., 2013, 2019; Galindon et al., 2016).

Thus, the results of our SDM analyses support the hypothesis that the small distribution ranges and high island endemicity of *R. lagascae*, *R. lobata* and *R. speciosa* cannot be explained by narrow environmental requirements of their host species (Pelser et al., 2018) and most probably also not by the absence of these hosts in areas with suitable environmental conditions because of other reasons. Instead, these *Rafflesia* species might be confined to their current distribution areas by their own environmental requirements and/or by having limited inter-island dispersal abilities. The former hypothesis was previously considered unlikely (Pelser et al., 2018), because *Rafflesia* species have been observed in both primary forests and those in various stages of regeneration, across wide elevational ranges, and on different substrates (Nais, 2001; Barcelona et al., 2009). Yet, here we show that the environmental needs of some *Rafflesia* (e.g., *R. speciosa*) may be an important limiting factor, although others may have less specific habitat requirements (e.g., *R. lobata*). Ancheta (2021) concluded in a previous SDM study of two other Philippine *Rafflesia* species (*R. consueloae* and *R. schadenbergiana*) that suitable habitat for these island endemics is found throughout the Philippines. It is thus possible that poor inter-island dispersal ability is the more common reason for the island endemicity of Philippine *Rafflesia*. However, we cannot exclude the possibility that *Rafflesia* species are absent from areas with suitable environmental conditions despite the presence of their host species because of other reasons than poor inter-island dispersal ability. Perhaps there are genotypes of *Tetrastigma* host species that a given *Rafflesia* species is unable to parasitize. However, there is currently no evidence that supports this alternative hypothesis. Instead, all three *Rafflesia* species studied here are able to parasitize *Tetrastigma* species belonging to different phylogenetic lineages (Pelser et al. 2016), indicating their ability to infect plants of quite diverse genotypes. Although we did not study this for *R. lobata*, we also did not find evidence of host-race formation in *R. lagascae* and *R. speciosa* (Pelser et al., 2017, 2018), which would have indicated that differences between different *Tetrastigma* genotypes are significant enough to drive local adaptation in *Rafflesia*.

### Conservation of Philippine *Rafflesia*, now and into the future

A better understanding of the distribution, habitat requirements, and dispersal abilities of species can result in more effective conservation management plans (Bellard et al., 2012; Guisan et al., 2013; Driscoll et al., 2014; Cai et al., 2022). This includes identifying areas that should be prioritized for formal protection (e.g., Spiers et al., 2018). Climate change may impact the size and location of suitable habitat in the future (e.g., Wilkening et al., 2019; Mancera and Lapuz, 2020). Understanding changes under different climate change scenarios can therefore inform management plans aimed at mitigating the impacts of climate change. This could consist of changing the boundaries of protected areas and establishing new protected areas to accommodate future changes to distribution areas of threatened species and to ensure or enhance connectivity among protected areas. Parks et al. (2023) showed that the Philippines is one of the countries that has some of the highest climate connectivity failures in Asia. This emphasizes the urgency to review its protected area network and the need for studies like ours to inform this work.

All Philippine *Rafflesia* species are of conservation concern (Pelser et al., 2011 onwards; DENR, 2017; Malabrigo et al., 2023). Many of them are only known from one or a handful disjunct populations and these sometimes appear to be composed of a single *Tetrastigma* host plant. Previous population genetic studies show strong genetic structure among populations of *R. lagascae* and *R. speciosa*, indicating low genetic connectivity among populations more than c. 20 km away from each other (Pelser et al., 2017, 2018). The Philippine tropical rainforest has been reduced to an estimated 3–6% of its original cover (Mittermeier et al., 1998; Ong et al., 2002) and the ongoing destruction and degradation of natural forest ecosystems in the Philippines are likely to increase distances between populations in this already fragmented landscape, lower their numbers, and reduce their sizes. Ultimately, this can result in a loss of genetic diversity through genetic drift and inbreeding (Ellstrand and Elam, 1993) and, consequently, populations that are less resilient and therefore more prone to extinction (Frankham, 2005). Species with patchy distributions and poor dispersal abilities, such as *Rafflesia*, are particularly vulnerable to this (Wang et al., 2015). SDM models under RCP 4.5 and RCP 8.5 indicate that climate change will exacerbate this threat for Philippine *Rafflesia*. All three *Rafflesia* species included in this study are predicted to lose suitable habitat on the islands on which they are currently found (Tables 2 and 3; Figs. 4–6), but *R. lagascae* might be most severely impacted, potentially losing as much as 14.3% of the area in which suitable habitat is currently found (Table 1). Our results suggest that *Rafflesia* populations at lower elevations might be particularly impacted, as climate change will see suitable habitat shifting to higher elevations (Appendix S6). It is presently unclear if populations will be able to track these elevation shifts. This will depend on their dispersal ability as well as the impact that climate change will have on habitats at local levels at these higher elevations. Because this is likely to vary from population to population, more detailed studies are required, such as those aimed at understanding changes to edaphic factors at local levels. Our data suggest that it might be important to consider such changes, because suitable areas for all three *Rafflesia* species under the current climate and the two climate change scenarios show small but statistically significant differences in edaphic variables, although it remains to be determined if these changes are also biologically significant and are likely to impact the potential for elevation shifts (Appendix S7). *Rafflesia* populations at lower elevations are also the populations that are likely to be the most prone to habitat destruction as a result of land conversion by expanding human population centers for residential and agricultural use, because they are typically close to places where people live and work (Hughes, 2017). Other *Rafflesia* species, both in the Philippines as in other countries, face similar challenges (e.g., Kusuma et al., 2022; Renjana et al., 2022; Malabrigo et al., 2023).

*Rafflesia* populations outside of protected areas are particularly vulnerable, now and in the future. For all three species and under each climate change scenario except for one (*R. speciosa* for RCP 4.5), the percentage of suitable area within the boundaries of present-day protected areas will reduce (Table 2). Although *R. lobata* is the species that is predicted to lose the least amount of suitable habitat in its native island range as a result of climate change (Table 1, Fig. 6a), none of its populations are located in protected areas (Appendix S8). Forest destruction, degradation and fragmentation as a result of logging, mining, farming, roading, new or expanding settlements, and other forms of encroachment thus appear to pose a greater risk to the survival of this species than climate change. About a third of the populations of *R. speciosa* are located in protected areas (Appendix S8). Most of these are at locations close to their boundaries and therefore more exposed to habitat degradation (Pelser et al., 2018). In addition, many *R. speciosa* populations are at sites that are at risk of losing suitable habitat in the future (Fig. 5a). One of the recommendations resulting from a conservation genetic study of *R. speciosa* (Pelser et al., 2018) was that the entire CPMR should be given protective status. The results of the present study provide further support for this recommendation and indicate that this would also be the best course of action to protect *R. lobata*. In Luzon, many *R. lagascae* populations are at the fringes of areas with suitable habitat and at or close to locations that are predicted to lose this habitat in the future (Fig. 4a). In addition, populations in the northern part of Luzon are all located outside of the few protected areas in this part of the country (Fig. 1). Because of this and the disjunct distribution of *R. lagascae* populations (Pelser et al., 2017), strategically-located new protected areas should be considered to expand the protected area network in Luzon and improve its connectivity.

## CONCLUSIONS

*Rafflesia* species are flagships of plant conservation. Recent studies improved our understanding of their species diversity, reproductive biology, propagation, environmental preferences, patterns of genetic diversity and connectivity, evolutionary relatedness, biogeographic history, and host preference and specificity in the Philippines (e.g., Barcelona et al., 2014; Galindon et al., 2016; Pelser et al., 2016, 2017, 2018, 2019; Molina et al., 2017; Lit, 2019; Tolod et al., 2020; Ancheta, 2021; Manuel and Hermocilla, 2021; Tobias et al., 2023). This SDM study of *R. lagascae, R. lobata* and *R. speciosa* built on these insights. Its results suggest that they are not rare because their *Tetrastigma* host species are rare. Instead, their island endemicity, small number of populations and disjunct distribution are most likely a result of poor inter-island dispersal abilities and/or specific environmental requirements, which appear to vary among the three species. Furthermore, all three species are projected to lose suitable habitat as a result of climate change, both within and outside protected areas on their native islands. However, some species are predicted to be more vulnerable than others. Additionally, the effectiveness of the current protected area network in the Philippines in meeting the conservation needs of these species varies among the three species, both presently and in the future. By understanding the similarities and differences among these three *Rafflesia* species, conservation strategies can be developed to protect each species as well as Philippine *Rafflesia* as a whole.

## ACKNOWLEDGMENTS

We are grateful to current and former staff of regional and local offices of the Department of Environment and Natural Resources (DENR), the Protected Area Management Board (PAMB) of Aurora Memorial National Park, Bataan Natural Park, Mt. Banahaw-San Cristobal Protected Landscape, Mt. Kanlaon Natural Park, Sibalom Natural Park, and Pantabangan-Carranglan Watershed Forest Reserve for facilitating the issuance of collecting and transport permits. We thank the local government units, communities, and field assistants and guides in Barbaza, Culasi, Pandan, San Remigio, Sibalom and Valderamma (Antique); Baler and Maria Aurora (Aurora); Mariveles (Bataan); Itogon (Benguet); Lal-o and Gattaran (Cagayan); Alcoy, Boljoon, Dalaguete, Cebu City, and Argao (Cebu); Leon, Igbaras, and Miag-ao (Iloilo); Bauang (La Union); Los Baños (Laguna); Dagami (Leyte); Dolores (Quezon); and Rodriguez (= Montalban) and Tanay (Rizal). For logistical support and assistance in the field, we are grateful to H. Alburo (Cebu Technological University-Argao), V. Salares (University of San Carlos), V. L. Dacumos (Nueva Ecija University of Science and Technology or NEUST), J. R. Callado and D. N. Tandang (National Museum of the Philippines), Aurora State College of Technology (ASCOT), L. R. Heaney and D. S. Balete (Field Museum, Chicago, IL, USA), Makiling Center for Mountain Ecosystem (MCME), M. A. O. Cajano (Museum of Natural History, University of the Philippines, Los Baños (UPMNH)). Thanks to our NGO partners: The Antique Outdoors (TAO), Cagayan Valley Partners in People Development (CAVAPPED) through P. A. Visorro, Conservation International, Philippines, Foundation for Philippine Environment (FPE), Philippines Biodiversity Conservation Foundation (PBCFI) through Lisa Paguntalan, and Tanggol Kalikasan, Quezon through J. Lim. D. J. V. Laborada and S. M. A. Ong (University of the Philippines Manila) assisted with or performed the preliminary species distribution modeling analyses for *Tetrastigma* species. Two anonymous reviewers generously volunteered their time and expertise to improve an earlier version of this manuscript. We value their kind and constructive feedback. This work was supported by the Marsden Fund Council from Government funding administered by the Royal Society of New Zealand [grant number UOC1102] and the Rufford Small Grant Foundation [grant number 20207-1].

## AUTHOR CONTRIBUTIONS

J. J. A. O., R. S. L., and P. B. P. conceptualized the research and methodology including data analysis and interpretation. P. B. P., J. F. B., and J. J. A. O. collected the data. P. B. P., J. J. A. O., R. S. L., and J. F. B. wrote and revised the manuscript. All authors gave their approval for the final version of the manuscript.

