## Appendix S1 for "What explains the high island endemicity of Philippine *Rafflesia*? A species distribution modeling analysis of three threatened parasitic plant species and their hosts"

Appendix S1. List of Maxent feature classes, regularization multipliers, and number of points used for modeling each species. L = Linear, Q = Quadratic; P = Product; H = Hinge; T = Threshold.

| **Species** | **Maxent FC** | **Regularization Multiplier** | **No. of Points** |
| --- | --- | --- | --- |
| *Rafflesia speciosa* | LQHPT | 1.5 | 40 |
| *Rafflesia lagascae* | L | 1.5 | 36 |
| *Rafflesia lobata* | H | 3 | 6 |
| *Tetrastigma* cf. *magnum* | LQH | 3 | 47 |
| *Tetrastigma harmandii* | LQ | 1 | 24 |
| *Tetrastigma loheri* | LQHPT | 1 | 79 |
| *Tetrastigma ellipticum* | LQH | 4.5 | 12 |
| *Tetrastigma* sp.A | LQHPT | 2.5 | 14 |
