## Appendix S2 for "What explains the high island endemicity of Philippine *Rafflesia*? A species distribution modeling analysis of three threatened parasitic plant species and their hosts"

Appendix S2. Average training Area Under the Curve (AUC) and True Skill Statistic (TSS) scores for the distribution models of each species.

| **Species** | **Average Training Area Under the Curve (AUC) Score** | **Average True Skill Statistic (TSS) Score** |
| --- | --- | --- |
| *Rafflesia speciosa* | 0.997 | 0.966 |
| *Rafflesia lobata* | 0.970 | 0.835 |
| *Rafflesia lagascae* | 0.972 | 0.840 |
| *Tetrastigma loheri* | 0.979 | 0.611 |
| *Tetrastigma sp. A* | 0.945 | 0.713 |
| *Tetrastigma ellipticum* | 0.876 | 0.477 |
| *Tetrastigma harmandii* | 0.930 | 0.515 |
| *Tetrastigma cf. magnum* | 0.963 | 0.661 |
