## Appendix S3 for "What explains the high island endemicity of Philippine *Rafflesia*? A species distribution modeling analysis of three threatened parasitic plant species and their hosts"

Appendix S3. Percent contribution values of each environmental variable for each individual SDM. Variables marked with (–) were not used in modeling the final distribution of that species. Values in bold were the highest for that species.

| **Environmental variable** | ***R. lagascae*** | ***R. speciosa*** | ***R. lobata*** | ***T. loheri*** | ***T. cf. magnum*** | ***T. ellipticum*** | ***T. sp. A*** | ***T. harmandii*** |
| --- | --- | --- | --- | --- | --- | --- | --- | --- |
| Mean Annual Temperature | **45.5** | 16.3 | 29.9 | **31.2** | **38.4** | **68.4** | **49.4** | 1.3 |
| Temperature Seasonality | 35.8 | 15.5 | – | 29.1 | 19.7 | 17.3 | 18.1 | 3.5 |
| Annual Precipitation | 0.1 | 19.7 | **70.1** | 7.9 | 5.6 | 3.5 | 13.8 | 24.3 |
| Precipitation Seasonality | 1.9 | **23.2** | – | 2.8 | 2.3 | 0.7 | 0 | 14.5 |
| Precipitation of the Coldest Quarter | 5.3 | 12.6 | – | 4.3 | 11.7 | 4.2 | 1.0 | 21.7 |
| Sand % | 0 | 10.8 | – | 3.0 | 7.4 | – | 4.7 | **29.3** |
| Silt % | 0.3 | 0.2 | – | 0.5 | 0 | 0 | – | 1.1 |
| Clay % | – | 0.9 | – | 1.4 | 13.6 | – | 0.7 | 2.8 |
| Soil pH | 11.0 | 1 | – | 11 | 0.8 | 5.7 | 0.8 | 0.2 |
| Soil Cation-Exchange Capacity | 0.1 | – | – | 4.0 | 0.5 | – | – | 1.0 |
| Bulk Density | – | – | – | 4.7 | – | 0.1 | 11.4 | 0.3 |
