## Appendix S4 for "What explains the high island endemicity of Philippine *Rafflesia*? A species distribution modeling analysis of three threatened parasitic plant species and their hosts"

Appendix S4. Gain-loss statistics for *Rafflesia* species relative to that of the islands from which they have been recorded (native island ranges) and the entire Philippines indicating changes in their amount of suitable habitat under the RCP 4.5 and RCP 8.5 climate scenarios in SDMs using *Rafflesia* occurrence points only (A1).

|  |  | **Native island ranges** | | **Entire Philippines** | |
| --- | --- | --- | --- | --- | --- |
|  | % | **RCP 4.5** | **RCP 8.5** | **RCP 4.5** | **RCP 8.5** |
| *Rafflesia lagascae* | **Gains** | 0.49 | 0.49 | 0.31 | 0.55 |
|  | **Losses** | 9.78 | 14.82 | 4.18 | 6.05 |
|  | **Stable: Suitable** | 22.30 | 17.26 | 8.70 | 6.83 |
|  | **Stable: Unsuitable** | 67.43 | 67.43 | 86.81 | 86.57 |
| *R. speciosa* | **Gains** | 1.21 | 1.14 | 0.27 | 0.29 |
|  | **Losses** | 2.15 | 4.65 | 0.21 | 0.44 |
|  | **Stable: Suitable** | 5.92 | 3.42 | 0.51 | 0.28 |
|  | **Stable: Unsuitable** | 90.73 | 90.80 | 99.00 | 98.99 |
| *R. lobata* | **Gains** | 0.00 | 0.00 | 0.33 | 0.40 |
|  | **Losses** | 0.92 | 1.48 | 0.28 | 0.88 |
|  | **Stable: Suitable** | 6.79 | 6.23 | 5.59 | 4.98 |
|  | **Stable: Unsuitable** | 92.29 | 92.29 | 93.80 | 93.74 |
