## Appendix S5 for "What explains the high island endemicity of Philippine *Rafflesia*? A species distribution modeling analysis of three threatened parasitic plant species and their hosts"

Appendix S5. Mean elevation (m) ± standard deviation (m) of *Rafflesia* species suitable habitat under different climate scenarios. Superscripts indicate significant differences between scenarios for each species.

|  | **Climate Scenarios** | | |
| --- | --- | --- | --- |
| **Species** | **Current** | **RCP 4.5** | **RCP 8.5** |
| *Rafflesia lagascae* | 785.46^a^ ± 423.77 | 965.88^a^ ± 397.75 | 1076.86^a^ ± 381.00 |
| *Rafflesia speciosa* | 674.59^b^ ± 310.26 | 827.60^b^ ± 296.62 | 1023.23^b^ ± 269.28 |
| *Rafflesia lobata* | 941.21^c^ ± 274.48 | 976.60^b^ ± 266.77 | 997.71^c^ ± 263.14 |
