## Appendix S8 for "What explains the high island endemicity of Philippine *Rafflesia*? A species distribution modeling analysis of three threatened parasitic plant species and their hosts"

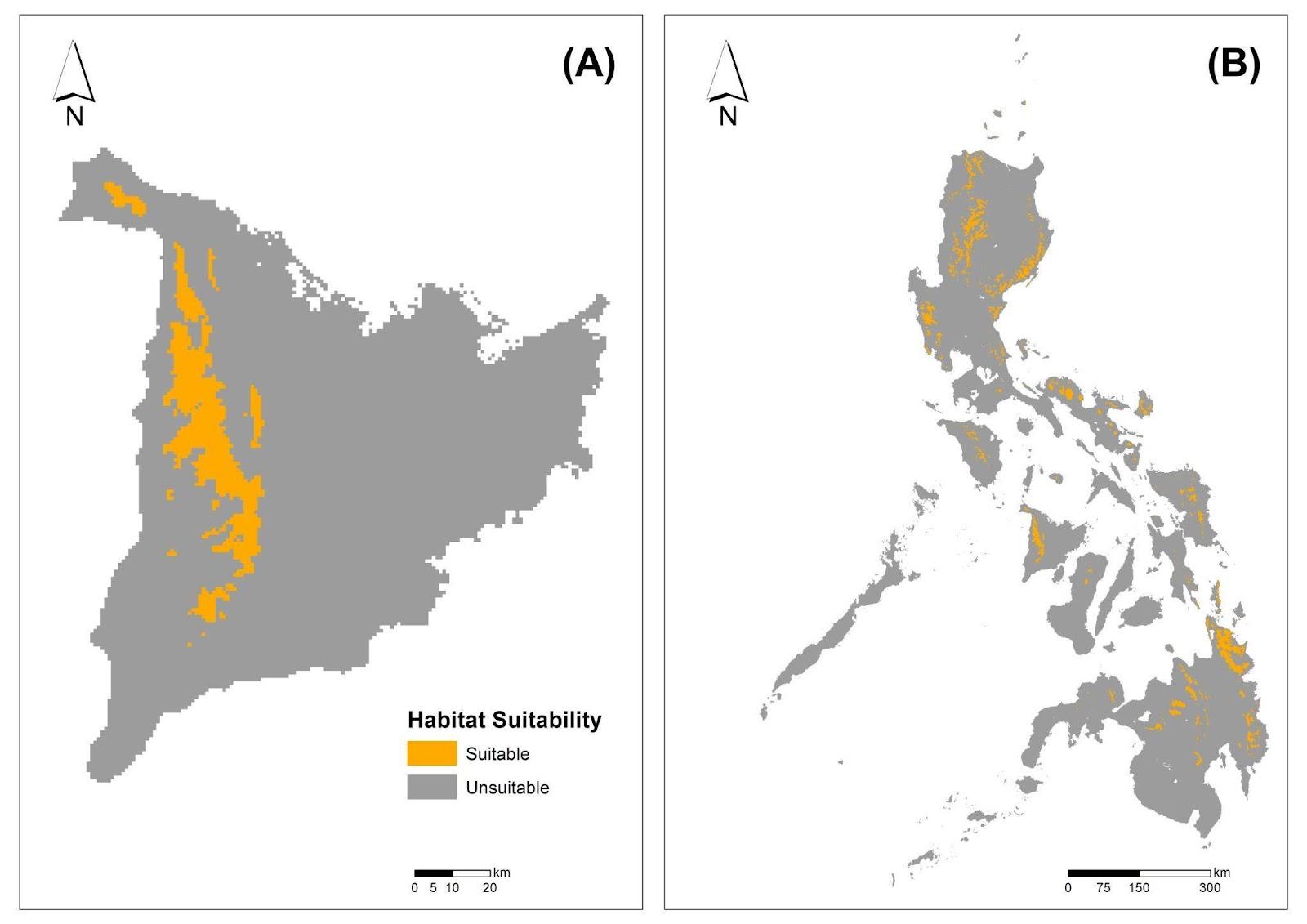

Appendix S8. Distribution of suitable habitat of *Rafflesia lobata* on Panay, the only island from which it is known (A) and the entire Philippines (B) under current climate conditions based on *R.* *lobata* SDMs masked with *Tetrastigma* habitat SDMs (A2 approach).
